# Implications of soil nitrogen enhancement on the yield performance of soybean (*Glycine max*) in cadmium-polluted soil

**DOI:** 10.1101/2021.11.22.469589

**Authors:** Beckley Ikhajiagbe, Matthew Chidozie Ogwu, Ivhuobe Izuapa Omoayena

## Abstract

The growth, development and yield of important crop plants like soybean *(Glycine max)* are constantly under threat by continuous inputs of cadmium in the biosphere as a result of various industrial activities. This study investigated the level to which, addition of nitrogen fertilization can enhance plant survival, growth and yield development in a cadmium-polluted. **T**hree accessions of *Glycine max* (TGm - 1, TGm - 2 and TGm - 3) were sown in a 12 mg/kg-cadmium polluted, which was thereafter amended with urea (FU), ammonia (FA), and ammonium nitrate (FN) singly and in combinations of equal proportions. A non-fertilized Cd-polluted soil and a general control constituted the negative and positive controls. **R**esults showed that although N application did not enhance yield dispositions of soybean in Cd-polluted soil, significant impact on vegetative development was noteworthy. Compared to yield of control plants, cadmium pollution imposed a 26.1 % reduction in per plant yield in TGm-1, compared 1.71 % in TGm-3. Generally, addition of nitrogenous fertilizer further suppressed crop yield by as much as 80 % in plants sown in cadmium-polluted soil. However, application of ammonia fertilizer to TGm-2 improved its yield performances in the cadmium-polluted soil.

## INTRODUCTION

Socio-economic crops serve as means of export to countries around the world and Soybean (*Glycine max*) provides about 64% of the world’s oilseed meal supply as well as 28 % of total vegetable oil production (USDA, 2000). Soybean is a major economic crop. In some developing economies, like Nigeria, particularly in rural regions, soybean symbolizes the unsurpassed source of non-meat protein accessible for enhancing the dietetic value of most traditional foods (Seralathan and Thirumaran, 1998). The preference of soybean as a better alternative for meat proteins is predicated on the fact that the seeds contain high protein levels, and its amino acid composition is approximate to the composition of animal proteins (Sikorski, 2007).

Cultivation of soybean seeds from farms, the availability of essential mineral elements, including favorable environmental conditions must be guaranteed; among such mineral elements is nitrogen. Among mineral elements, nitrogen is required by plants in large amounts. It acts as a constituent of several plant cell components, including amino acids as well as nucleic acids.

Thus, the deficiency of nitrogen inhibits plant growth rapidly. When such a deficiency persists, most species display chlorosis (yellowing of the leaves). This is often seen in older leaves close to the base which ultimately fall off the plant. However, younger leaves may not display these symptoms initially because nitrogen can be mobilized from older leaves.

In plants, the toxicity of cadmium is often associated with imbalances in several macro - and micronutrient levels as well as growth inhibition. In addition, the presence of cadmium in plants results in several physiological alterations and these affects both carbohydrate and nitrogen metabolism as reported by Chaffei *et al*. (2003). Cadmium also induces modifications in the membrane functionality and Howlett and Avery (1997) reported this process to occur by the changes triggered in the lipid composition of membranes. It has also been shown that cadmium-induced toxicity also involves some other senescence-like processes. A good example is the involvement of oxidative stress, which is likely mediated by hydrogen peroxide. This eventually leads to gradual increases in the activities of some antioxidant enzymes, namely peroxidase and catalase, accompanied by increased lipid peroxidation (Sandalio *et al*., 2001).

Since vegetal species such as soybean are able to accumulate large quantities of cadmium in their edible parts, there is a serious health risk if they are grown on soils polluted with cadmium. Serious diseases have been reported by Oliveira *et al*. (1999) to be associated with the ingestion of food and/or water polluted with cadmium even when the cadmium concentration is considered to be in low amount. Thus, due to the high dependence on cash crops such as soybean (*Glycine max*) to man, anima and the generation of income, this research was therefore carried out to investigate if, the addition of different nitrogen fertilizers to cadmium polluted soil will enhance the growth and performance of *Glycine max*.

## MATERIALS AND METHOD

### Source of seeds and soil

The seeds of three different accessions of *glycine max* (TGm 1, TGm 2 and TGm 3) used in this study were obtained from IITA Ibadan, Nigeria. Top soil was used for this study while Nitrogen sources used were ammonia, urea and ammonium nitrate. Field experiment was conducted at the University botanical garden.

### Pollution of soil and addition of nitrogen sources

Pollution was done by adding 0.2g (×3 ESV) of cadmium in bags containing 20kg of top soil and mixing properly with a hand trowel. A total of 81 bags (27 bags per accession) were used. Only one concentration of the stipulated quantity of cadmium was diluted in 1.5litres of water, thoroughly mixed and poured into the bags. The same gram of cadmium served as a treatment for all the bags excluding the control for all accessions which also lacked the addition of a nitrogen source.

The polluted bags were allowed to seat for a period of 4 days before been fertilized. The nitrogen sources of ammonia, urea and ammonium nitrate were allotted FU, FA and FN respectively. FU for ammonia, FA for urea and FN for ammonium nitrate. Treatments of FU, FA, FN, FU+FA, FU+FN, FA +FN, FU+FA+FN were administered to bags - 9grams of each of ammonia, urea and ammonium nitrate (FU, FA and FN), 6grams each of the fertilizers ( FU+ FA, FU + FN, FA + FN) and 3grams each (FU + FA + FN), excluding the control which was left unfertilized and heavy metal free. Another treatment was unfertilized but polluted. Each treatment was carried out on all three seed accessions, including triplicates. Soil physio – chemical analysis (see Table 1), following the method of AOAC (2002) was done before pollution took place.

**Table 1:** Physical and chemical properties of soil before application of nitrogen treatments.

| Parameters | Soil |
| --- | --- |
| pH | 5.27 |
| Electrical conductivity ($\mu\text{s}/\text{cm}$) | 301.21 |
| Total organic carbon (%) | 0.49 |
| Total Nitrogen (%) | 0.18 |
| Exchangeable acidity (meq/100 g soil) | 0.22 |
| Na (meq/100 g soil) | 10.90 |
| K (meq/100 g soil) | 1.48 |
| Ca (meq/100 g soil) | 14.32 |
| Mg (meq/100 g soil) | 12.01 |
| $\text{NO}_2$ (mg/kg) | 16.43 |
| $\text{NO}_3$ (mg/kg) | 30.01 |
| Clay (%) | 5.13 |
| Silt (%) | 7.06 |
| Sand (%) | 87.81 |
| Fe (mg/kg)* | 1011.92 |

*Soil is ferruginous*

### Planting of seeds and germination studies

Four seeds of *Glycine max* per bag were sown in all treatments, including the control. During a 12-week period, morphological characters were observed and recorded.

#### Growth and performance studies

the growth parameters were determined, alongside data analysis using SPSS version 20 software while Statistical analysis was conducted under a 95% confidence limit.

## RESULTS AND DISCUSSION

Table 1, the pH of the soil gave a value of 5.27; with an electric conductivity of 301.21μs^−1^cm. Total soil nitrogen was 0.18%. It is however necessary to state that the soil before use was ferruginous because the iron content was 1011.92mg^−1^kg. The soil also had better infiltration properties having a textural sand content of 87.81%.

Figure 1 shows the effects of different nitrogen application on plant height of matured 12weeks old soybean plants. TGm-1 for the control, recorded a plant height of 29.80 cm compared to TGm-1 in the urea (URR) enhanced cadmium contaminated soil, which had a plant height of 31.70 cm. However, the lowest height for TGm-1 was obtained in the cadmium contaminated soil enhanced with a combination of ammonia and ammonium nitrate (AAN) - 17.70 cm. Generally, TGm-2 in the control was higher in height compared to the other plants in cadmium contaminated soil. This is actually expected as the presence of cadmium would have caused a significant reduction in the plant height.

**Fig 1:**
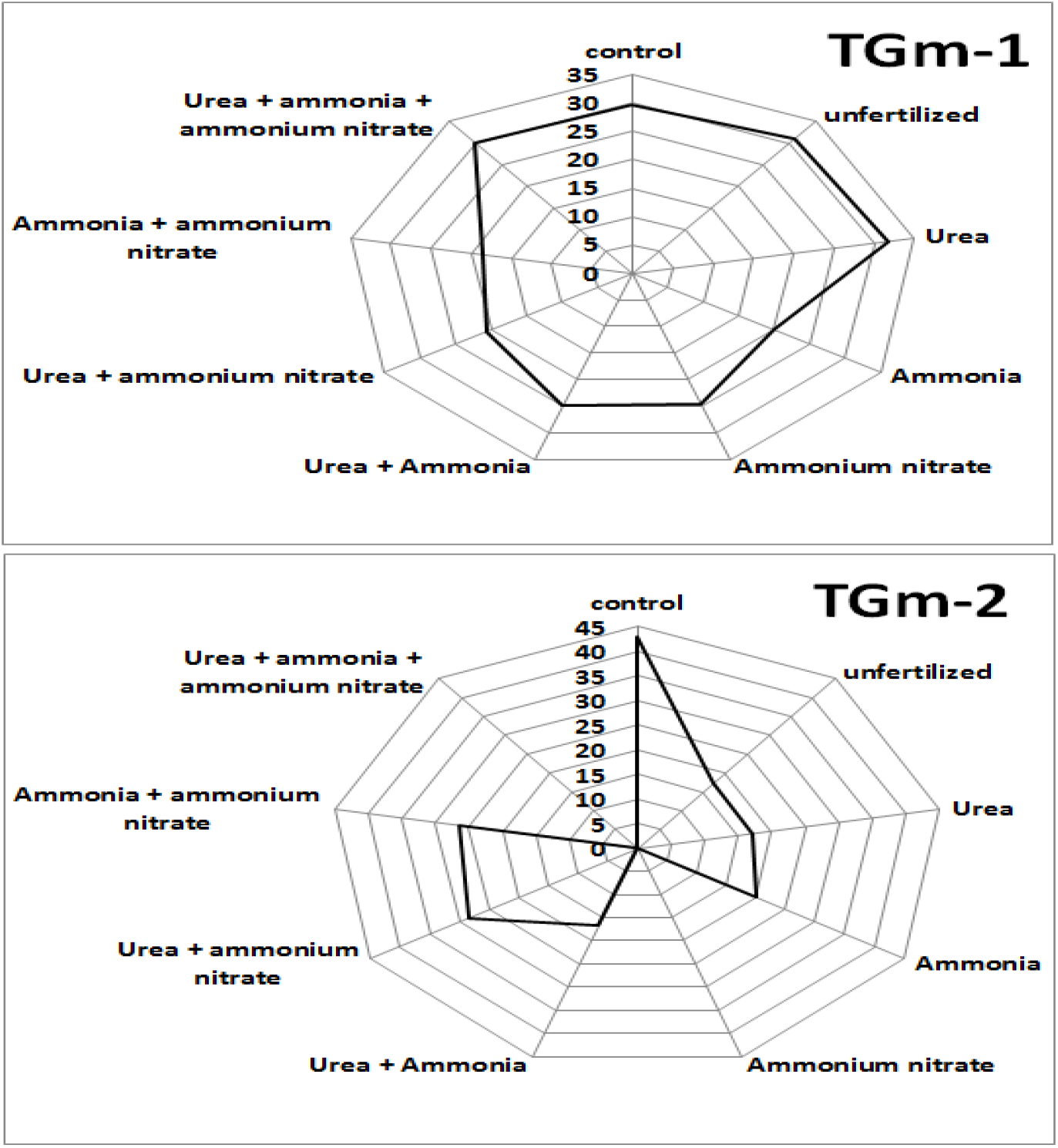
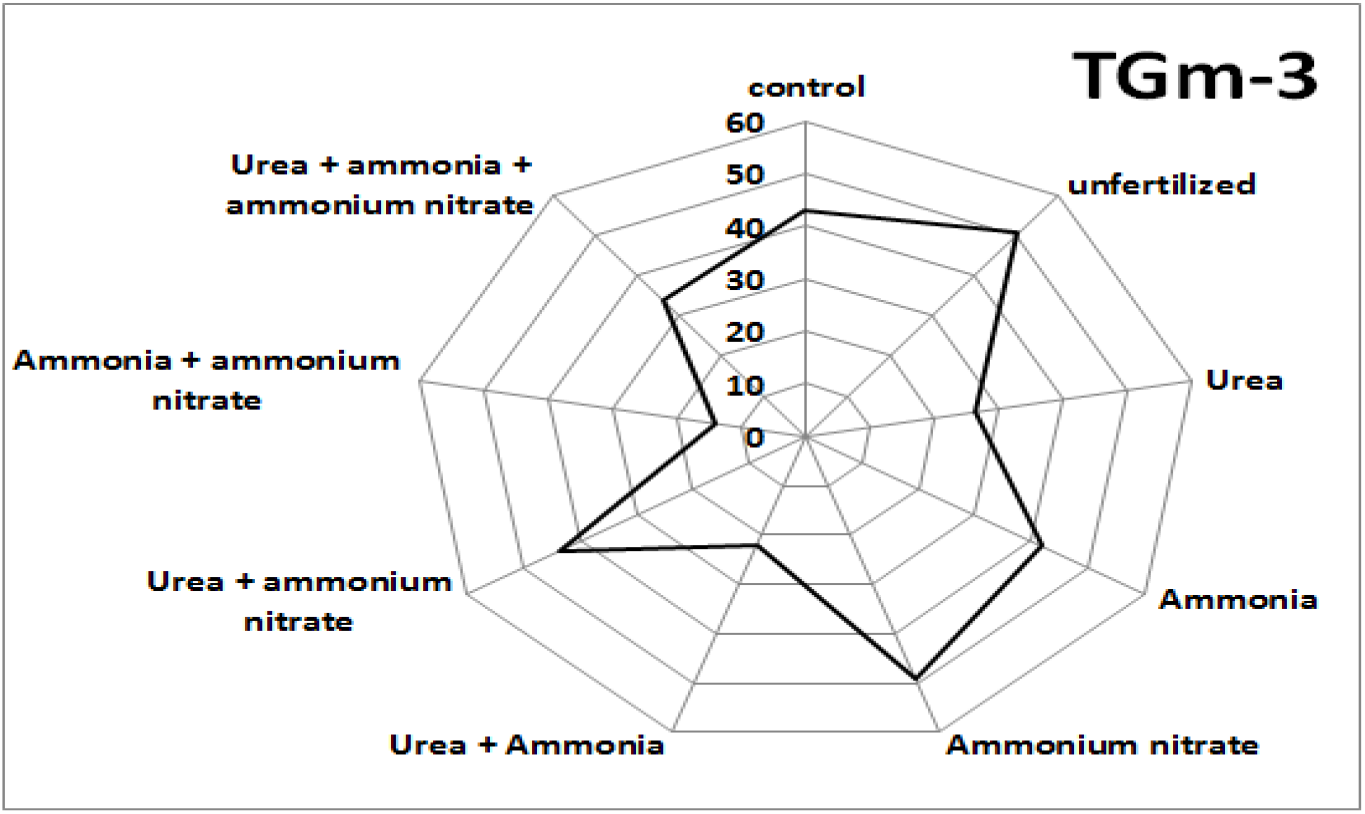
Effects of differing nitrogen applications on plant height of mature soybean.

However, upon addition of nitrogen to the soil, there was an increase in the plant height of TGm-2. For example - in the unfertilized soil - with the addition of ammonia (AMM), the plant height increased to 17.4cm. This height was further increased to 20 cm as well as to 28.50 cm upon the addition of urea and ammonium nitrate (UAN) to the soil. TGm-3 of the unfertilized had the highest height (50.50 cm) compared to all other treatments in this cultivar. However, with the addition of AAN a decrease in height was noted (14.0 cm). The addition of cadmium did not necessarily affect growth in TGm-3 as plants in the polluted soil fared as good as the control. AAN positively influenced the plant’s growth as the plant reached a height of 49.0cm which was higher than that of the control (43.0 cm). In TGm-3, the control as well as plants in soils enhanced with the nitrogen sources generally experienced greater heights when compared to TGm-1 and TGm-2; with TGm-2 having the least height.

(see Figure 2).

**Figure 2:**
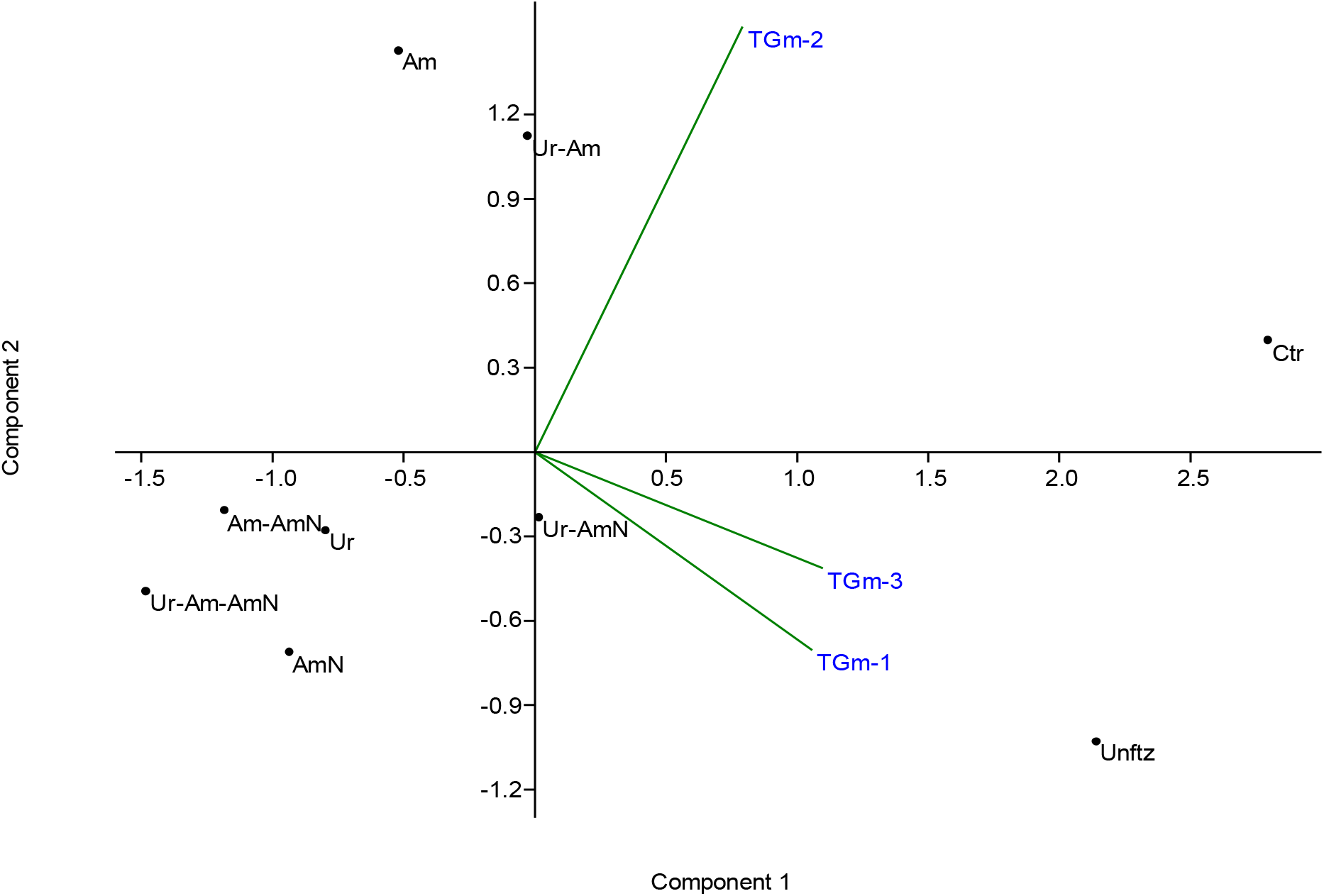
Principal component plot showing relationship existing between the nitrogen nutrient applications and per plant yield dispositions of the 3 test soybean accessions.

The effects of the different applications of nitrogen in the patterns of stem branching have been reported in table 2. In the control plant there were two primary stem branches in TGm-1. However, for TGm-2 and TGm-3, there were 6 primary branches. Results showed that cadmium application enhanced primary stem branching in soybean plants. For example, in the unfertilized cadmium contaminated soil, there was an increase in stem branching from 2 to 4 for TGm-1, as well as 6 to 8 in TGm-2. The results remained so when nitrogen was applied to the soil. However, in some cases there was reduction upon application of nitrogen. For example, the urea, ammonia and ammonium nitrate (ALL) enhanced cadmium contaminated soil stem branching. In TGm-1, primary stem branching reduced to 3 as against 8 and also reduced to 5 for TGm-3 as against 9.

**Table 2:**
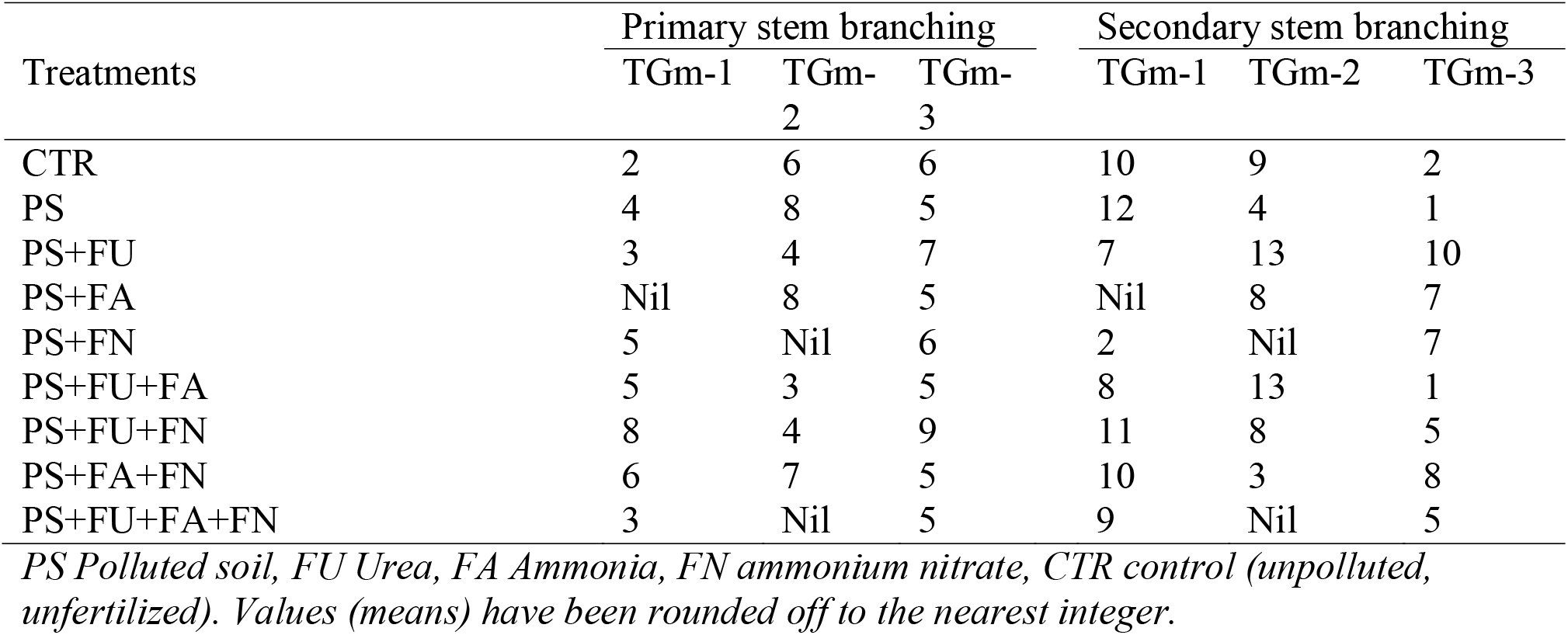
Effects of differing nitrogen applications on stem branching characteristics of mature soybean.

For secondary stem branching, plant responded differently. For example, there was an increase in secondary stem branching for TGm-1 from 10 in the control to 12 in the unfertilized cadmium polluted soil. There was significant reduction for TGm-2 from 9 to 4 in the unfertilized cadmium polluted soil. It is however important to note that application of nitrogen in the cadmium contaminated soil enhanced generally production of stem branching, although there were a few exceptions; for example, in the application of urea and ammonia (URA) for TGm-3.

In comparing the number of primary stem branches between all three soybean cultivars, TGm-3 generally had the highest amount of primary stem branches, with the exclusion of a few treatments (unfertilized, AMM and AAN). Conversely, in comparing the number of secondary stem branches between the cultivars, TGm-1 generally had the highest number of secondary stem branches.

Differences in vegetative growth between N source treatments in both the plants were apparent by full bloom when the number of primary and secondary stem branches was greater than in un-augmented soils. As seen in Table 2, a combination of UAN had the highest number of primary stem branches and the application of U had the highest number of secondary stem branches.

It was also noticed generally that there was a presence of pigmentation in the stem, branch, petiole and peduncle in both cadmium contaminated soils and control, as well as in the fertilized cadmium contaminated soil (Table 3). However, plant responded differently with regards to the intensity of the pigmentation. Although pigmentation was not extensive in the various plant parts accessed, it however ranged from slight to moderate. According to the Methuen color chart code as provided by IITA, leaf color ranged from dark green to green.

**Table 3:**
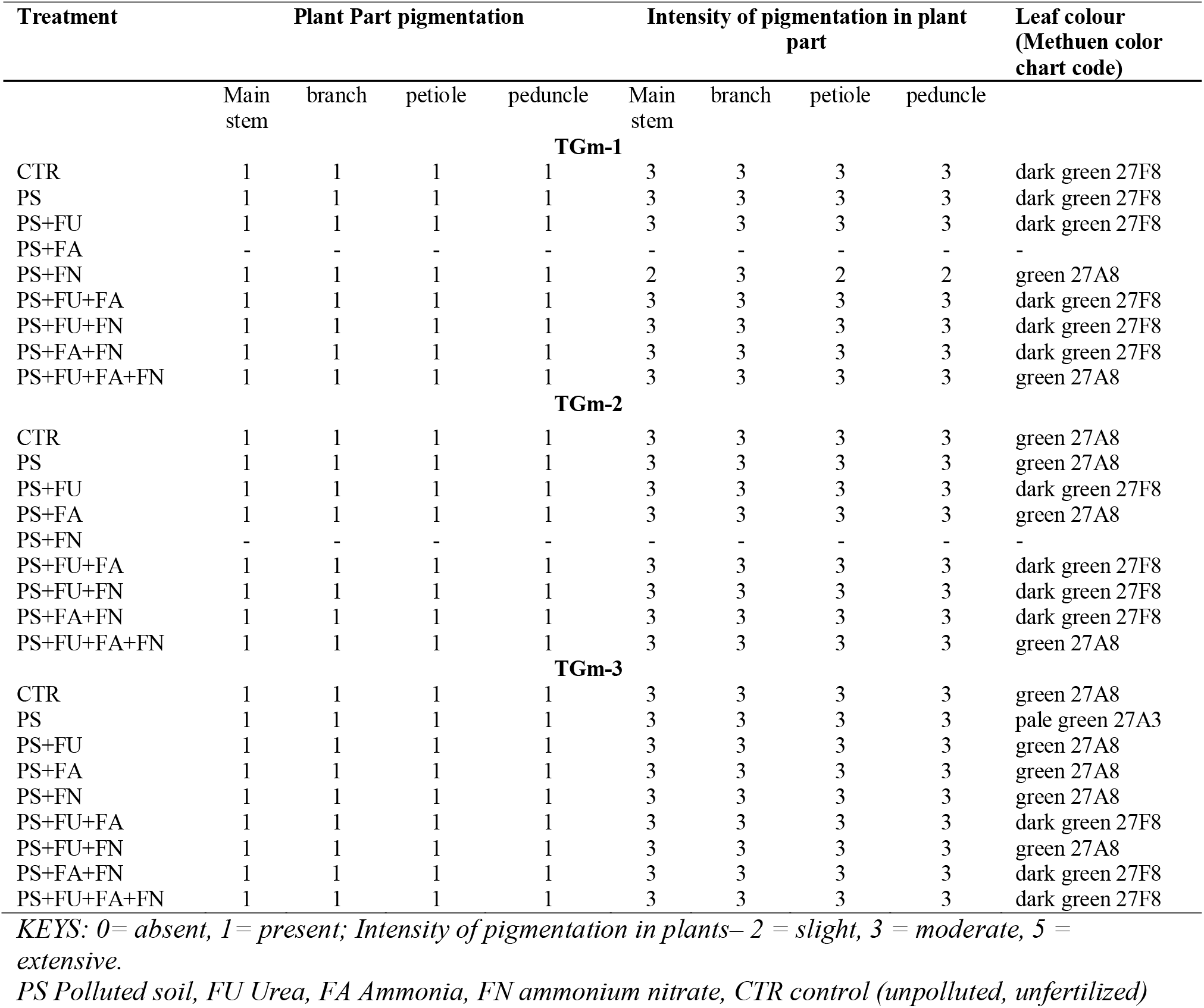
changes in Intensity of pigmentation in plant parts upon exposure to differing nitrogen sources.

| Treatment | Plant Part pigmentation |  |  |  | Intensity of pigmentation in plant part |  |  |  | Leaf colour (Methuen color chart code) |
| --- | --- | --- | --- | --- | --- | --- | --- | --- | --- |
|  | Main stem | branch | petiole | peduncle | Main stem | branch | petiole | peduncle |  |
| TGm-1 |  |  |  |  |  |  |  |  |  |
| CTR | 1 | 1 | 1 | 1 | 3 | 3 | 3 | 3 | dark green 27F8 |
| PS | 1 | 1 | 1 | 1 | 3 | 3 | 3 | 3 | dark green 27F8 |
| PS+FU | 1 | 1 | 1 | 1 | 3 | 3 | 3 | 3 | dark green 27F8 |
| PS+FA | - | - | - | - | - | - | - | - | - |
| PS+FN | 1 | 1 | 1 | 1 | 2 | 3 | 2 | 2 | green 27A8 |
| PS+FU+FA | 1 | 1 | 1 | 1 | 3 | 3 | 3 | 3 | dark green 27F8 |
| PS+FU+FN | 1 | 1 | 1 | 1 | 3 | 3 | 3 | 3 | dark green 27F8 |
| PS+FA+FN | 1 | 1 | 1 | 1 | 3 | 3 | 3 | 3 | dark green 27F8 |
| PS+FU+FA+FN | 1 | 1 | 1 | 1 | 3 | 3 | 3 | 3 | green 27A8 |
| TGm-2 |  |  |  |  |  |  |  |  |  |
| CTR | 1 | 1 | 1 | 1 | 3 | 3 | 3 | 3 | green 27A8 |
| PS | 1 | 1 | 1 | 1 | 3 | 3 | 3 | 3 | green 27A8 |
| PS+FU | 1 | 1 | 1 | 1 | 3 | 3 | 3 | 3 | dark green 27F8 |
| PS+FA | 1 | 1 | 1 | 1 | 3 | 3 | 3 | 3 | green 27A8 |
| PS+FN | - | - | - | - | - | - | - | - | - |
| PS+FU+FA | 1 | 1 | 1 | 1 | 3 | 3 | 3 | 3 | dark green 27F8 |
| PS+FU+FN | 1 | 1 | 1 | 1 | 3 | 3 | 3 | 3 | dark green 27F8 |
| PS+FA+FN | 1 | 1 | 1 | 1 | 3 | 3 | 3 | 3 | dark green 27F8 |
| PS+FU+FA+FN | 1 | 1 | 1 | 1 | 3 | 3 | 3 | 3 | green 27A8 |
| TGm-3 |  |  |  |  |  |  |  |  |  |
| CTR | 1 | 1 | 1 | 1 | 3 | 3 | 3 | 3 | green 27A8 |
| PS | 1 | 1 | 1 | 1 | 3 | 3 | 3 | 3 | pale green 27A3 |
| PS+FU | 1 | 1 | 1 | 1 | 3 | 3 | 3 | 3 | green 27A8 |
| PS+FA | 1 | 1 | 1 | 1 | 3 | 3 | 3 | 3 | green 27A8 |
| PS+FN | 1 | 1 | 1 | 1 | 3 | 3 | 3 | 3 | green 27A8 |
| PS+FU+FA | 1 | 1 | 1 | 1 | 3 | 3 | 3 | 3 | dark green 27F8 |
| PS+FU+FN | 1 | 1 | 1 | 1 | 3 | 3 | 3 | 3 | green 27A8 |
| PS+FA+FN | 1 | 1 | 1 | 1 | 3 | 3 | 3 | 3 | dark green 27F8 |
| PS+FU+FA+FN | 1 | 1 | 1 | 1 | 3 | 3 | 3 | 3 | dark green 27F8 |

**KEYS:** 0= absent, 1= present; Intensity of pigmentation in plants– 2 = slight, 3 = moderate, 5 = extensive.
PS Polluted soil, FU Urea, FA Ammonia, FN ammonium nitrate, CTR control (unpolluted, unfertilized)

In TGm-1, pigmentation was present in the plant’s main stem, branch, petiole and peduncle. This was also the same for TGm-2 and TGm-3, as these cultivars were noticed to also be pigmented. Intensity of pigmentation in plant parts of TGm-1 were moderate, with the exception of plants grown in soil enhanced with ammonium nitrate (ANN). Slight pigmentation was noted in the main stem, petiole and peduncle of plants enhanced with ANN.

Dark green leaf coloration was noticed in TGm-1; a slight deviation from dark-green to green was influenced by the application of ANN and ALL. Prominent leaf color of TGm-3 was darkgreen and green, with the exception of the leaves of the unfertilized plants having pale green coloration.

It should be noted that application of cadmium neither affected the pigmentation in plant part, intensity of plant part pigmentation nor the leaf coloration as there was no difference in the control and most of other polluted treatments among all three cultivars.

Intensity of pigmentation in plant parts with the application of differing nitrogen sources had noticeably similar characteristics; pigmentation in main stem, branch, petiole and peduncle was present, intensity did not vary in these parts and leaf color ranged from green to dark green. The above also prove true for the control and plants grown in unfertilized soils. This result indicates that cadmium pollution did not influence photosynthetic rates and pathways. This is in line with works that have proven that plants are able to optimize the allocation of N in order to present a balance between the calvin cycle and light harvesting (cholorophyll) capabilities (Warren and Adams, 2001).

The percentage occurrence of foliar chlorosis was taken into note (Table 4). Results showed that the pattern of occurrence of foliar chlorosis was similar among all 3 accessions (p = 0.634). Within each accession, soil amendment with the various nitrogen applications significantly impacted on the level of development of chlorosis; for example, in TGm-2, percentage chlorosis in Cd-contaminated plants significantly reduced from 53.10 % in the unfertilized Cd-polluted soil to between 2.20 % – 19.10 % in the nitrogen-fertilized Cd-polluted soils (p = 0.026). Generally, there was no significant difference in the occurrence of foliar necrosis throughout the study in TGm-1 (p = 0.429), TGm-2 (p = 0.226) and TGm-3 (p = 0.302) respectively. As observed, the incident of cadmium pollution did not cause any form of foliar folding in soybean compared to the control; however, upon elevation of soil nitrogen, significant increases in foliar folding was reported in all 3 accessions as high as 80 % (Table 5).

**Table 4:** Effects of differing nitrogen applications on some physical characteristics of soybean monitored to post flowering, before pod production.

| Treatments | Chlorosis (%) | Necrosis (%) | Leaf folding (%) | Senesced leaves (%) |
| --- | --- | --- | --- | --- |
| **TGm-1** |  |  |  |  |
| CTR | 18.50 | 3.90 | 0 | 10.20 |
| PS | 14.64 | 0 | 0 | 13.0 |
| PS+FU | 6.30 | 0 | 66.30 | 15.80 |
| PS+FA | 0 | 0 | 55.10 | 16.30 |
| PS+FN | 17.80 | 0 | 42.50 | 19.20 |
| PS+FU+FA | 3.40 | 0 | 36.0 | 13.50 |
| PS+FU+FN | 0 | 0 | 51.20 | 19.50 |
| PS+FA+FN | 5.70 | 0 | 48.60 | 20.0 |
| PS+FU+FA+FN | 7.50 | 0 | 33.80 | 13.40 |
| *p-value (within gp.)* | **0.048** | **0.429** | **0.023** | **0.182** |
| **TGm-2** |  |  |  |  |
| CTR | 19.10 | 2.70 | 0 | 2.70 |
| PS | 53.10 | 3.30 | 0 | 68.80 |
| PS+FU | 0 | Nil | 0 | 87.50 |
| PS+FA | 2.20 | 0 | 40.0 | 5.60 |
| PS+FN | 16.70 | 0 | 0 | 0 |
| PS+FU+FA | 4.60 | 0 | 13.60 | 63.60 |
| PS+FU+FN | 0 | 0 | 41.70 | 25.0 |
| PS+FA+FN | 15.40 | 0 | 53.9 | 7.70 |
| PS+FU+FA+FN | 12.3 | 3.14 | 8.51 | 3.8 |
| *p-value (within gp.)* | **0.026** | **0.226** | **0.021** | **0.018** |
| **TGm-3** |  |  |  |  |
| CTR | 3.50 | 0 | 0 | 8.40 |
| PS | 3.80 | 0.60 | 0 | 7.90 |
| PS+FU | 8.70 | 0 | 14.30 | 4.0 |
| PS+FA | 1.30 | 0 | 1.30 | 13.0 |
| PS+FN | 21.40 | 0 | 15.40 | 11.70 |
| PS+FU+FA | 3.20 | 0 | 51.0 | 15.60 |
| PS+FU+FN | 35.70 | 0 | 21.0 | 3.60 |
| PS+FA+FN | 0 | 0 | 82.80 | 13.80 |
| PS+FU+FA+FN | 1.10 | 0 | 23.90 | 17.0 |
| *p-value (within gp.)* | **0.049** | **0.302** | **0.009** | **0.056** |
| p-value (between gps.) | 0.634 | 0.442 | 0.213 | 0.138 |
| *PS Polluted soil, FU Urea, FA Ammonia, FN ammonium nitrate, CTR control (unpolluted, unfertilized)* |  |  |  |  |

**Table 5:** Effects of differing nitrogen applications on floral characteristics of soybean.

|  | Days to 50%<br>peduncle<br>initiation<br>(days) | Days to<br>50%<br>flower bud<br>initiation<br>(days) | Flowers<br>/peduncle | Flowering<br>duration<br>(days) |
| --- | --- | --- | --- | --- |
| **TGm-1** |  |  |  |  |
| CTR | 36 | 40 | 28 | 70 |
| PS | 33 | 38 | 33 | 67 |
| PS+FU | 35 | 37 | 35 | 63 |
| PS+FA | 32 | 36 | 32 | 67 |
| PS+FN | 33 | 38 | 35 | 66 |
| PS+FU+FA | 36 | 36 | 33 | 67 |
| PS+FU+FN | 35 | 36 | 29 | 71 |
| PS+FA+FN | 32 | 37 | 36 | 64 |
| PS+FU+FA+FN | 33 | 35 | 33 | 68 |
| *p-value (within gp.)* | **0.098** | **0.028** | **0.274** | **0.011** |
| **TGm-2** |  |  |  |  |
| CTR | 35 | 50 | 23 | 68 |
| PS | 35 | 47 | 26 | 65 |
| PS+FU | 34 | 45 | 32 | 59 |
| PS+FA | 33 | 44 | 33 | 58 |
| PS+FN | 33 | 47 | 28 | 61 |
| PS+FU+FA | 34 | 48 | 32 | 60 |
| PS+FU+FN | 35 | 46 | 33 | 63 |
| PS+FA+FN | 32 | 46 | 33 | 65 |
| PS+FU+FA+FN | 33 | 48 |  |  |
| *p-value (within gp.)* | **0.104** | **0.052** | **0.261** | **0.005** |
| **TGm-3** |  |  |  |  |
| CTR | 34 | 49 | 31 | 69 |
| PS | 33 | 48 | 33 | 67 |
| PS+FU | 35 | 48 | 28 | 59 |
| PS+FA | 32 | 43 | 35 | 62 |
| PS+FN | 33 | 45 | 29 | 67 |
| PS+FU+FA | 32 | 47 | 28 | 65 |
| PS+FU+FN | 32 | 44 | 31 | 65 |
| PS+FA+FN | 33 | 45 | 30 | 67 |
| PS+FU+FA+FN | 32 | 47 | 36 | 63 |
| *p-value (within gp.)* | **0.143** | **0.016** | **0.242** | **0.014** |
| *p-value (between gps.)* | 0.121 | 0.000 | 0.238 | 0.014 |

*PS Polluted soil, FU Urea, FA Ammonia, FN ammonium nitrate, CTR control (unpolluted, unfertilized)*
*Mean values have been presented to the nearest integer*

When compared to cereals and grasses, legume crops are less tolerant to cadmium toxicity and legumes encounter strong inhibition of biomass production due to cadmium (Inouhe *et al*., 1994). Cadmium toxicity also impairs the physiological as well as biochemical processes in plants namely photosynthesis, mineral uptakes and water relations (Lopez–chuken and young, 2010; Gill *et al*., 2012). In addition, Gratao *et al*. (2009) and Gallego *et al*. (2012) also reported disruption of membrane function and composition. These findings lead to the suggestion that leaf folding which was only present in cadmium polluted soils (Table 4) was as a result in the impairment in the physiological and biochemical processes in the test plant.

This suggestion proved true due to the fact that the control had no leaf fold, although results of Table 4 also suggest a relationship between nitrogen augmentation in soils and leaf folding. Data suggest that the changes observed in nitrogen metabolism in Cd-stressed plants are similar to changes observed during natural senescence (Masclaux *et al*., 2000) as the main function of leaf senescence is the recycling of nutrients. These findings are in line with the percentage senesced leaves noted in Table 4. Much significant difference was not noticed between the control and the test plant grown in cadmium polluted soil. Although, unfertilized plants and URR enhanced plants of TGm-2 where noted to have the highest percentage of senesced leaves.

In Table 5, presentation of floral characteristics of the test plant in Cd-polluted soil after exposure to soil nitrogen elevation was similar in all 3 plant accessions. No significant differences were noticed in the number of days taken for plant to achieve 50 % peduncle initiation (32 – 36 days). However, plants showed significant differences in the number of days required to attain 50 % flower bud initiation. Similarly, there were differences in flower duration was obtained (p = 0.014); values ranged from 59 – 71 days.

In all three plant cultivars (TGm-1, TGm-2 and TGm-3), controls were noted to have the longest days to 50 % flower bud initiation. However, pollution with cadmium and enhancement with nitrogen reduced the duration of flower bud initiation; for example, with the addition of ALL in TGm-1 50 % of flower buds were initiated in 35 days while the control took 40 days.

The addition of cadmium is also noted to have influenced flowering duration in TGm-1. In comparison to the control (70days), all other treatments had shorter days in flowering; with an exception of UAN which had a longer flowering duration of 71days. In TGm-2 and TGm-3, increased levels of nitrogen affected the duration of flowering; the control had longer days of flowering when compared to other treatments.

In Table 6, cadmium application and the differing nitrogen applications had no effect on pod characteristics of mature soybean (*Glycine max*). No significant difference was noted in the seed cavity ridges on pods and pod dehiscence; control and all other treatments of the three accessions had seed cavity ridges present and pods were non – shattering.

**Table 6:** Effects of differing nitrogen applications on pod characteristics of mature soybean pods.

| Treatments | Seed cavity ridges on pods |  |  | Pod dehiscence |  |  |
| --- | --- | --- | --- | --- | --- | --- |
|  | TGm-1 | TGm-2 | TGm-3 | TGm-1 | TGm-2 | TGm-3 |
| CTR | 1 | 1 | 1 | 0 | 0 | 0 |
| PS | 1 | 1 | 1 | 0 | 0 | 0 |
| PS+FU | 1 | 1 | 1 | 0 | 0 | 0 |
| PS+FA | Nil | 1 | 1 | Nil | 0 | 0 |
| PS+FN | 1 | Nil | 1 | 0 | Nil | 0 |
| PS+FU+FA | 1 | 1 | 1 | 0 | 0 | 0 |
| PS+FU+FN | 1 | 1 | 1 | 0 | 0 | 0 |
| PS+FA+FN | 1 | 1 | 1 | 0 | 0 | 0 |
| PS+FU+FA+FN | 1 | Nil | 1 | 0 | Nil | 0 |

*PS Polluted soil, FU Urea, FA Ammonia, FN ammonium nitrate, CTR control (unpolluted, unfertilized)*
*Pod dehiscence: 0 = non – shattering , 1 = shattering. Seed cavity ridges on pods: 0 = absent, 1 = present*

In Table 7, all cultivars were noted to have similar seed characteristics; absence of split testa, seed shapes predominantly oval, no variegation in seeds and brown testa with continuous narrow black shape around the hilum was noticed. Testa texture was mostly smooth in TGm-1and seed brilliance was medium, with the exception of AAN. The addition of AAN produced seeds which were round. Only the unfertilized plants of TGm-2 produced wrinkled seeds. However, the wrinkling did not affect the brilliance of its seeds. Seeds produced with the application of UAN were matt.

Generally, the addition of cadmium and application of the differing nitrogen applications had no significant influence on seed characteristics.

**Table 7:** Effects of differing nitrogen applications on seed characteristics of harvested soybean seeds, according to seed description by IITA.

| Treatments | Splitting of testa | Seed shapes | Testa texture | Testa color variegation | Basal color of variegated seeds | Eye color of white seeds | Eye color pattern | Brilliance of seeds |
| --- | --- | --- | --- | --- | --- | --- | --- | --- |
| **TGm-1** |  |  |  |  |  |  |  |  |
| CTR | 0 | 2 | 2 | 0 | 0 | 0 | 1 | 2 |
| PS | 0 | 2 | 1 | 0 | 0 | 0 | 1 | 2 |
| PS+FU | 0 | 2 | 2 | 0 | 0 | 0 | 1 | 2 |
| PS+FA | Nil | Nil | Nil | Nil | Nil | Nil | Nil | Nil |
| PS+FN | 0 | 2 | 2 | 0 | 0 | 0 | 1 | 2 |
| PS+FU+FA | 0 | 2 | 1 | 0 | 0 | 0 | 1 | 2 |
| PS+FU+FN | 0 | 2 | 1 | 0 | 0 | 0 | 1 | 2 |
| PS+FA+FN | 0 | 1 | 1 | 0 | 0 | 0 | 1 | 1 |
| PS+FU+FA+FN | 0 | 2 | 2 | 0 | 0 | 0 | 1 | 2 |
| **TGm-2** |  |  |  |  |  |  |  |  |
| CTR | 0 | 2 | 1 | 0 | 0 | 0 | 1 | 2 |
| PS | 0 | 2 | 3 | 0 | 0 | 0 | 1 | 2 |
| PS+FU | 0 | 2 | 2 | 0 | 0 | 0 | 1 | 2 |
| PS+FA | 0 | 2 | 1 | 0 | 0 | 0 | 1 | 2 |
| PS+FN | Nil | Nil | Nil | Nil | Nil | Nil | Nil | Nil |
| PS+FU+FA | 0 | 2 | 2 | 0 | 0 | 0 | 1 | 2 |
| PS+FU+FN | 0 | 2 | 1 | 0 | 0 | 0 | 1 | 1 |
| PS+FA+FN | 0 | 2 | 1 | 0 | 0 | 0 | 1 | 2 |
| PS+FU+FA+FN | Nil | Nil | Nil | Nil | Nil | Nil | Nil | Nil |
| **TGm-3** |  |  |  |  |  |  |  |  |
| CTR | 0 | 2 | 1 | 0 | 0 | 0 | 1 | 2 |
| PS | 0 | 2 | 1 | 0 | 0 | 0 | 1 | 1 |
| PS+FU | 0 | 2 | 1 | 0 | 0 | 0 | 1 | 1 |
| PS+FA | 0 | 2 | 2 | 0 | 0 | 0 | 1 | 2 |
| PS+FN | 0 | 2 | 1 | 0 | 0 | 2 | 1 | 2 |
| PS+FU+FA | 0 | 2 | 2 | 0 | 0 | 0 | 1 | 2 |
| PS+FU+FN | 0 | 2 | 1 | 0 | 0 | 0 | 1 | 2 |
| PS+FA+FN | 0 | 2 | 1 | 0 | 0 | 0 | 1 | 3 |
| PS+FU+FA+FN | 0 | 2 | 2 | 0 | 0 | 2 | 1 | 1 |

PS Polluted soil, FU Urea, FA Ammonia, FN ammonium nitrate, CTR control (unpolluted, unfertilized)
Splitting of testa - 0 = absent, 1 = present (testa splits to expose the cotyledons). Seed shapes - 1 = round/ globular, 2 = oval, 3 = oblong, 4 = rhomboid.
Testa texture, scored on the seeds by touching the testa surface as: 1 = smooth, 2 = rough, 3 = wrinkled (folds on the testa)
Testa colour variegation - 0 = absent, 1 = present
Basal colour of variegated seeds - 0 = non – variegated seeds, 1 = cream, 2 = brown, 3 = black
Eye colour of white seeds (color around the hilum of white seeds) - 0 = non – white seeds, 1 = clean (no color around the hilum), 2 = brown, 3 = black
Eye color pattern - 1 = brown testa with continuous narrow black stripe around the hilum, 2 = brown testa with dark brown fork-like eye pattern, 3 = white/grey testa with incision-like eye pattern, 4 = brown testa with dark brown incision-like pattern below and parallel to the hilum, 5 = white testa with reddish brown vase-like eye, 6 = white testa with black vase-like eye
Brilliance of seeds - 1 = matt, 2 = medium, 3 = shiny

Table 8 shows the effects on yield of soybean with the differing nitrogen applications. In TGm-1, the unfertilized plants produced pods weighting 33.8g; this was noted to be the largest weight. The differing nitrogen applications affected the weight of pods as these were greatly reduced when compared to pods produced by non-nitrogen enhanced plants; for example, the control weighed 35.1g, pods from plants enhanced with ANN weighted 6.6g and those of ALL weighted 4.93g.

**Table 8:** Effects of differing nitrogen applications on yield outcome of soybean.

| Treatments | Weight of pods per plant (g) |  |  | Number of pods* |  |  | Seeds/pods* |  |  | Yield per plant (g) |  |  |
| --- | --- | --- | --- | --- | --- | --- | --- | --- | --- | --- | --- | --- |
|  | TGm-1 | TGm-2 | TGm-3 | TGm-1 | TGm-2 | TGm-3 | TGm-1 | TGm-2 | TGm-3 | TGm-1 | TGm-2 | TGm-3 |
| CTR | 35.1 | 18.8 | 18.5 | 45 | 20.7 | 23 | 4 | 3 | 4 | 25.3 | 12.47 | 11.7 |
| PS | 33.8 | 7.2 | 17.7 | 30 | 11 | 21 | 3 | 3 | 2 | 18.7 | 5.4 | 11.5 |
| PS+FU | 5.3 | 3.57 | 14.7 | 11.7 | 3.3 | 16 | 3 | 4 | 3 | 3.17 | 2.05 | 5.57 |
| PS+FA | Nil | 11.5 | 9.93 | Nil | 17.7 | 11 | nil | 3 | 3 | Nil | 9.6 | 4.03 |
| PS+FN | 6.6 | Nil | 10.9 | 4.33 | Nil | 18 | 4 | Nil | 2 | 3.6 | Nil | 5.8 |
| PS+FU+FA | 7.07 | 12.1 | 14.1 | 7 | 18 | 18 | 3 | 3 | 4 | 4.07 | 9.4 | 5.4 |
| PS+FU+FN | 13.9 | 5.63 | 13 | 20.3 | 3.67 | 17 | 3 | 3 | 4 | 12.4 | 4.27 | 5.7 |
| PS+FA+FN | 8.4 | 7.07 | 5.23 | 11 | 5.33 | 4 | 3 | 3 | 3 | 7.07 | 2.03 | 2.33 |
| PS+FU+FA+FN | 4.93 | Nil | 8.33 | 7 | Nil | 9 | 3 | nil | 3 | 3.03 | Nil | 3.13 |
| p-value (within gps.) | 0.001 | 0.038 | 0.046 | 0.000 | 0.027 | 0.041 | 0.423 | 0.413 | 0.293 | 0.001 | 0.020 | 0.027 |
| p-value (between gps.) | 0.318 |  |  | 0.307 |  |  | 0.462 |  |  | 0.264 |  |  |

*\*presented to the nearest whole number*
*PS Polluted soil, FU Urea, FA Ammonia, FN ammonium nitrate, CTR control (unpolluted, unfertilized)*

There was a significant drop in the weight of pods of the unfertilized plants in TGm-2 (7.2g), however, pod weight increased with the application of AMM and URA; weighing 11.5g and 12.1g respectively. Cadmium is suspected to have influenced these weights, although the different nitrogen applications resulted in a slight significant increase of pod weights. TGm-3 responded differently when compared to the other accessions. The application of UAN increased weight of pods from 9.93g in AMM to 14.1g.URR enhanced soils, produced pods weighing 14.7g, this was in close range to the control (18.3g).

URA and ALL in TGm-1 both had 7 pods, which also is the least number of pods. As to be expected, the weight of pods per plant was proportional to the number of pods produced. Plants in unpolluted soils had the largest number of pods in TGm-1. Cadmium significantly affected the number of pods produce; for example, control with approximately 21 pods, unfertilized produced 30 pods and UAN produced 7 pods. It is important to note that the differing nitrogen applications did not particularly increase the number of pods. In TGm-2 and TGm-3, the control was also noted to have the higher number of pods, although with the application of nitrogen, the effect of cadmium was significantly reduced.

An average of 3 seeds per pod was noted in TGm-1, TGm-2 and TGm-3. However, the application of AAN increased seed number to 4 in TGm-1. Unfertilized of TGm-3 had 2 seeds per pod and no significant effect with the application of nitrogen was noticed in the number of seeds produced per pod.

There was however significant difference in yield of TGm-1 as unfertilized plants and control had the highest yield (18.7 g and 25.3 g) respectively. With the application of nitrogen, plant yield significantly dropped; for example, 3.17 g with the application of URR and 4.07 g with the application of URA. Plant yield in TGm-2 dropped to 5.4 g when unfertilized but increased with the addition of AMM. In TGm-3 the reverse was the case; addition of nitrogen reduced yield of plants, with 2.33 g been the lowest yield when AAN was applied.

The seed yield in terms of weight of pods per plant, number of pods, and yield per plant was similar among the different nitrogen applications as noted in Table 9. High yields were noticed in the control and unfertilized plants. Although the different combinations of nitrogen fertilizers were supposed to have enhanced yield of the test plant, these influenced the yield of plants negatively. The application of 6 g of urea to the soil decreased the weight of pods from 33.8 g in the fertilized to 5.3 g in TGm-1, and in TGm-3 pod weight decreased from 17.7 g to 14.7 g. A combined application of 3g each of urea and ammonium nitrate only increased the weight of pods by 1.77 g, and the application of 2 g each of urea, ammonia and ammonium nitrate further decreased plant yield to 4.93 g. Yield per plant was highest in the unfertilized. With the application of 6 g of ammonia, yield decreased to 4.03 g from 11.5 g. Further enhancement with 3 g each of ammonia and ammonium nitrate reduced yield to 2.33 g.

**Table 9:** Percentage yield reduction and recovery from repressed growth.

| Treatments | Overall yield reduction (%) |  |  | Recovery from metal-induced growth repression (%) |  |  |
| --- | --- | --- | --- | --- | --- | --- |
|  | TGm-1 | TGm-2 | TGm-3 | TGm-1 | TGm-2 | TGm-3 |
| CTR | RV | RV | RV | - | - | - |
| PS | 26.1 | 56.7 | 1.71 | RV | RV | RV |
| PS+FU | 87.5 | 83.6 | 52.4 | -83.1 | -62.3 | -51.6 |
| PS+FA | Nil | 23.2 | 65.6 | Nil | 77.8 | -65.1 |
| PS+FN | 85.8 | Nil | 50.4 | -80.7 | Nil | -49.6 |
| PS+FU+FA | 83.9 | 24.6 | 53.8 | -78.2 | 74.1 | -53.4 |
| PS+FU+FN | 51.3 | 65.8 | 51.3 | -33.7 | -20.9 | -50.4 |
| PS+FA+FN | 72.1 | 83.7 | 80.1 | -62.2 | -62.4 | -79.7 |
| PS+FU+FA+FN | 88.1 | Nil | 73.2 | -83.8 | Nil | -72.8 |

*Negative values represent reductions. RV reference value.*

Compared to yield of control plants, there was a 26.1 % reduction in per plant yield in TGm-1 sown in unfertilized polluted soil (PS), compared to 87.5 yield reduction when the plant was further exposed to urea (Table 10). The lowest growth repression due to Cd pollution was obtained in TGm-3 (1.71 % reduction). Generally, addition of nitrogenous fertilizer further suppressed crop yield by as much as 80 %. However, application of ammonia as well as a combination of urea and ammonia enhanced yield dispositions of the cadmium-impacted plants by 77.8 % and 74.1 % respectively (Table 9). Consequently, application of ammonia fertilizer to TGm-2 improved its yield performances in a cadmium-polluted soil.

**Table 10:** Results for determination of nitrate nitrogen, ammonium nitrogen, and total nitrogen in leaf and root samples of the test plant during flowering.

|  | Leaves |  |  | Roots |  |  |
| --- | --- | --- | --- | --- | --- | --- |
|  | Nitrate<br>nitrogen<br>(ppm) | Ammonia<br>nitrogen<br>(ppm) | Total<br>nitrogen<br>(%) | Nitrate<br>nitrogen(p<br>pm) | Ammonia<br>nitrogen<br>(ppm) | Total<br>nitrogen<br>(%) |
| **TGm-1** |  |  |  |  |  |  |
| CTR | 1356.6 | 39.6 | 0.7 | 1627.0 | 31.8 | 0.5 |
| PS | 1978.4 | 48.8 | 3.1 | 1407.1 | 98.0 | 3.2 |
| PS+FU | 1624.2 | 55.3 | 1.3 | 2968.0 | 42.7 | 3.8 |
| PS+FA | 1634.5 | 42.4 | 4.2 | 1715.8 | 8.5 | 4.3 |
| PS+FN | 1536.5 | 75.4 | 5.3 | 1372.8 | 16.5 | 3.4 |
| PS+FU+FA | 1490.2 | 43.5 | 3.7 | 1831.3 | 18.6 | 3.3 |
| PS+FU+FN | 1933.4 | 44.7 | 2.4 | 1519.1 | 23.1 | 0.6 |
| PS+FA+FN | 1704.6 | 80.2 | 5.2 | 2455.2 | 30.3 | 1.3 |
| PS+FU+FA+FN | 1506.0 | 102.1 | 3.9 | 2005.3 | 22.8 | 1.7 |
| *p-value (within gp.)* | **0.523** | **0.099** | **0.048** | **0.086** | **0.367** | **0.411** |
| **TGm-2** |  |  |  |  |  |  |
| CTR | 1445.7 | 89.6 | 0.7 | 1138.1 | 38.5 | 1.1 |
| PS | 2891.5 | 21.9 | 1.6 | 1731.0 | 35.2 | 1.1 |
| PS+FU | 1623.3 | 33.5 | 2.5 | 1692.7 | 17.3 | 1.5 |
| PS+FA | 2012.2 | 32.3 | 2.6 | 2104.9 | 130.5 | 6.9 |
| PS+FN | 1012.8 | 61.0 | 5.2 | 1255.8 | 5.4 | 4.2 |
| PS+FU+FA | 1810.1 | 63.6 | 3.0 | 1962.8 | 14.4 | 1.5 |
| PS+FU+FN | 1119.2 | 53.4 | 3.4 | 1581.3 | 5.3 | 4.3 |
| PS+FA+FN | 1565.8 | 36.5 | 3.4 | 1989.1 | 26.3 | 2.5 |
| PS+FU+FA+FN | 1940.7 | 36.5 | 4.2 | 2033.4 | 21.5 | 2.2 |
| *p-value (within gp.)* | **0.637** | **0.202** | **0.042** | **0.093** | **0.041** | **0.396** |
| **TGm-3** |  |  |  |  |  |  |
| CTR | 2082.9 | 391.8 | 3.4 | 3416.6 | 12.7 | 2.0 |
| PS | 1003.0 | 58.3 | 5.3 | 1770.1 | 22.9 | 1.1 |
| PS+FU | 2833.4 | 70.4 | 5.0 | 3085.5 | 15.8 | 2.7 |
| PS+FA | 1901.2 | 72.9 | 5.3 | 2321.2 | 31.8 | 4.7 |
| PS+FN | 1169.2 | 56.4 | 5.3 | 1638.1 | 28.1 | 3.3 |
| PS+FU+FA | 1615.8 | 46.5 | 7.5 | 1721.0 | 26.4 | 4.3 |
| PS+FU+FN | 1682.6 | 50.3 | 4.7 | 1672.7 | 15.3 | 3.6 |
| PS+FA+FN | 1810.7 | 75.5 | 5.6 | 2102.3 | 53.3 | 4.5 |
| PS+FU+FA+FN | 2011.3 | 66.3 | 5.2 | 3194.9 | 60.0 | 5.0 |
| *p-value (within gp.)* | **0.523** | **0.099** | **0.067** | **0.040** | **0.101** | **0.380** |
| p-value (between gps.) | 0.782 | 0.234 | 0.003 | 0.071 | 0.935 | 0.426 |

*PS Polluted soil, FU Urea, FA Ammonia, FN ammonium nitrate, CTR control (unpolluted, unfertilized)*

As presented in Figure 2, an attempt was made to show relationship between the nitrogen nutrient application and yield dispositions of the three test plants. The yield of TGm-2 was most closely influenced by URA, whereas those of TGm-3 and TGm-1 were most likely influenced by UAN combinations.

A dendogram for all treatment applications have been presented (figure 3). The hierarchical cluster presented herein was an attempt to compare which treatments had close or near – similar effects on the plants. As presented, there were two major groups, the first was that which comprised of the control and the unfertilized soil. The other group tied up the unfertilized soils; a clear indication that the effect of nitrogen application in the soil was distinct (figure 3). Within the nitrogen enhanced plants, the most closely related were urea and ammonia nitrate, however the combination of UAN was a stand-alone.

**Figure 3:**
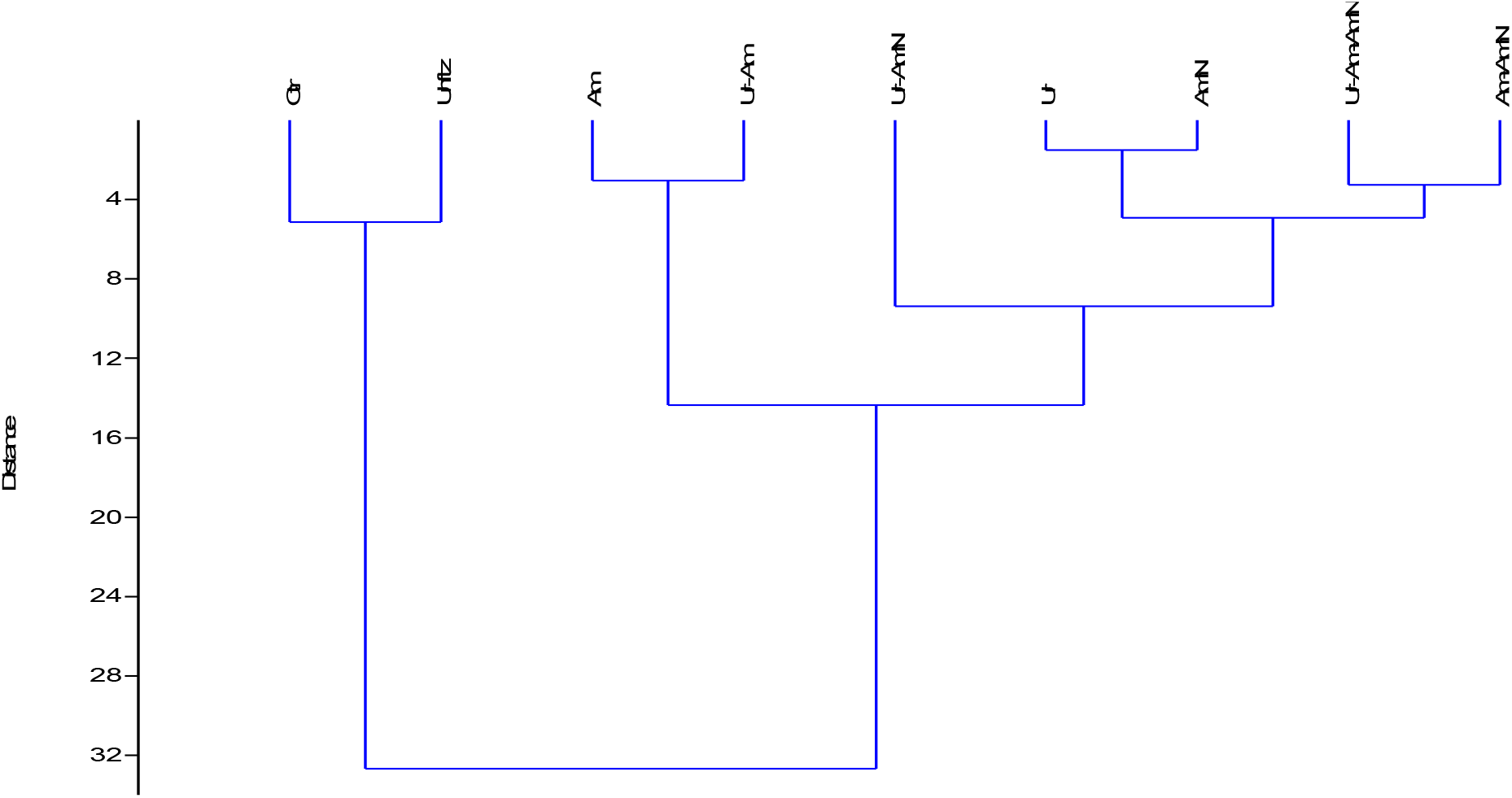
Hierarchical cluster dendrogram showing relationship existing between the nitrogen nutrient applications and per plant yield dispositions of the 3 test soybean accessions.

The determination of nitrate nitrogen (NN), ammonium nitrogen (AN) and total nitrogen (TN) in leaves and roots sample of the test plant during flowering was noted in Table 10. In leaves of TGm-1 analyzed, no significant difference was noted in NN and AN content. However, percentage TN increased with the application of nitrogen. Although TN content of unfertilized leaves was moderately high (3.1 %), it could however not be compared to leaves of plants enhanced with ANN (5.3 %) and AAN (5.2 %). The control had the lowest total leaf nitrogen content of 0.7 %. In the roots, NN content was highest (2968.0 ppm) when URR was applied and was lowest when AAN was applied. Unfertilized roots analyzed were noted to have 98.0 ppm of AN, but with the separate application of AAN, AN content reduced to 8.5ppm. Percentage TN content in roots was lowest in the control (0.5 %) and highest with the application of AAN. This is also in alignment with analysis carried out on leaves of TGm-1.

NN content of TGm-2’s leaves analyzed was noted to be highest in the unfertilized (2891.5 ppm). However, there was no notable significant difference with the addition of nitrogen. AN content was highest at 89.6 ppm for control, although addition of AAN and UAN increased AN levels to 61.0 ppm and 63.0 ppm respectively. As noticed in TGm-1, TGm-2 also had reduced total nitrogen content in the control. The addition of nitrogen is noted to have influenced the total nitrogen content of TGm-2. No significant difference was noted when roots were analyzed for NN and TN bur AN was high in the control and unfertilized when AAN was applied.

In TGm-3, NN was highest with the application of URR (2832.4 ppm) and lowest in unfertilized leaves. The control had a high amount of AN (391.8 ppm) when compared to other leaves of plants exposed to cadmium; for example, the enhancement of soils with ALL and URR produced leaves with an AN content of 66.3 ppm and 70.4 ppm respectively. Cadmium had no significant effect on leaves as TN content was noted to vary slightly vary from one treatment to the next (URR-5.0 %, URA – 7.5 %, and AAN – 5.6 %); although the control had a TN content of 3.4 %. The application of nitrogen is noticed to positively influence the TN content in leaves of TGm-3. NN content in the roots was highest in the control. The enhancement of soils with URR and ALL also influence this. Roots in soils to which URR was applied had NN content of 3085.5 ppm and those in ALL enhanced soils contained 3194.0 ppm. AN content was highest in roots to which ALL was applied (60.0 ppm). TN content increased in roots with the different nitrogen applications; from 4.7 % with the application of AMM to 5.0 % with the application of ALL. Roots of the unfertilized soil as noted to have the lowest TN content of 1.1 %.

It has been shown that enzymes of N metabolism are differently affected by Cd stress (Chugh *et al*., 1992). Nitrate reductase (NR) and nitrite reductase (NiR) activities are significantly decreased, leading to reduced nitrate assimilation by plants. The activities of the glutamine synthetase–glutamate synthase (GS/GOGAT) cycle, which operates in ammonium incorporation into carbon skeletons, are also affected (Gouia *et al*., 2000). These findings did not hold regarding the nitrate nitrogen content of the test plant. As noted in Table 11, the addition of UAN to soils increased the nitrate nitrogen content in leaves from 1356.6 ppm in the control, to 1933.4 ppm in TGm-1 and 1445.7 ppm in the control, to 2012.2 ppm with the application of ammonia in TGm-2. Total nitrogen content in leaves was 7.5 % with the application of URA and in roots this content increased with the application of AMM.

### Comparative Plant Response

In terms of yield, TGm-1 was noted to be the best cultivar with pods weighing 35.1 g, a total of 45 pods (4 seeds per pod) and a general yield of 25.4 g.TGm-3 had the most prominent nitrogen assimilatory response; in leaves, NN content was 2082.9ppm, AN content was 391.8 ppm and TN content was 3.4 %. While in the roots, NN content was 3416.6 ppm, AN was 12.7 ppm and TN was 2.0 %

TGm-3 was the most tolerant to pollution in terms of yield. In comparison to other cultivars, it had a total of 36 pods weighting 33.8 g and a yield of 18.7 g. In cadmium polluted soils, TGm-1 showed the most tolerance in terms of N – assimilatory response. Leaves of TGm-1 had NN content of 1978.4 ppm, AN content of 48.8 ppm and TN content of 3.1 %. NN present in its roots was 14072 ppm, AN - 98.0 ppm and TN – 3.2 %. TGm −1.

The combination of urea and ammonia nitrate (UAN) generally enhanced plant growth and development regardless of cadmium contaminated soils in which plants were grown. The application of this combination averaged the highest plant height in all three cultivars and was noted to increase the number of primary and secondary stem branches.

Ammonia enhanced the survival of the weakest soybean in terms of yield. It increased pod weight from 7.2 g to 11.5 g, number of pods from 11 to 19, and yield of plant from 5.4 g to 9.6 g. In terms of N – assimilatory response, survival of the weakest link was aided with the combined application of urea, ammonia and ammonium nitrate (ALL); for example, in leaves NN content of 1003.0ppm increased to 2011.3ppm and AN content from 58.3 ppm to 66.3 ppm. In roots, NN levels shot up to 3194.9 ppm from 1770.1 ppm, AN level had a significant rise of 60.0 ppm from 22.9 ppm, and TN content of 1.1 % increased to 5.0 %.

## CONCLUSION

The distribution and chemical behavior of environmental pollutants, such as Cadmium in plants of economic importance are of great significance in understanding, projecting, and limiting the effect of this harmful metal in the growth and development of plants such as soybean which produces more protein and oil than any other land plant. The enhancement of polluted soils with nitrogen sources is projected to limit the harmful effect of this metal in terms of growth and yield of crops. However, in this study N application did not enhance yield dispositions of soybean in Cadmium - polluted soil, but significant impact on vegetative development was noteworthy.

